# Adaptive and neutral processes that select on life course diversification

**DOI:** 10.1101/2022.09.06.506771

**Authors:** Ulrich Steiner, Shripad Tuljapurkar

## Abstract

Heterogeneity among individuals in fitness components is what selection acts upon. One of the fundamental questions in evolution is how such heterogeneity arises and is maintained. Fundamental evolutionary theories predict that selection acts against such heterogeneity but contrasting these predictions substantial non-genetic and non-environmental driven variability in phenotypes can be observed; not only under highly controlled lab conditions, but also in less controlled settings. Here we ask, by analysing structured population models across a large range of biological taxa, how selective forces act on the processes that generate variation among life courses. Our findings suggest that non-genetic, non-environmental driven variation is in general neither truly neutral, selected for, or selected against. Much variation exists among species and populations within species, with mean patterns suggesting close to neutral evolution of life course variability. Populations that show greater diversity of life courses do not show, in general, increased or decreased population growth rates, and selective forces acting on the processes of diversification seem not to generally increase or decrease life course variability. Approaches in quantitative genetics that identified similar lack of understanding of maintenance of variability might be extended to include non-genetic and non-environmental driven variability. We believe to be only at the beginning in understanding the evolution and maintenance of non-genetic non environmental variation.

## Introduction

Individuals in any population vary in their life courses, exemplified by differences in lifespan, reproduction, phenotypes, and functional traits [1–4]. Classical evolutionary theories, founded in seminal work by Fisher [5], Wright [6], and Haldane [7,8], explain such variation by genotypic variation, environmental variation, or their interaction. If constant environments exist over many generations, according to these theories, selection should erode genotypic variation by selecting for adaptive phenotypes and their associated genotypes. In population genetics terms additive genetic variation should erode, while neutral molecular variation maintains some genetic diversity without substantial phenotypic variation, if the phenotypes are selected upon [9,10]. In its consequence, in a constant environment, among individual variation in phenotypes and life courses should be reduced, if phenotypes are linked to fitness. These predictions are challenged by the observation that even isogenic individuals, originating from parental populations that have been exposed over many generations to highly controlled lab conditions, exhibit high levels of variation among individual life courses and phenotypes, also for phenotypes that directly link to fitness and that are under selection [11–13]. Similarly, in less controlled genetic and environmental conditions, environmental variation, genotypic variation, and their interaction only account for a small fraction of the total observed phenotypic variation of fitness components [14–17]. For systems where the decomposition among genotypic, environmental and other stochastic variation is challenging, due to lack of accurate data, similar amounts of total phenotypic variation can be observed as in more controlled systems [16,18,19]. The question arises how such high levels of phenotypic variation can be maintained as basic evolutionary theories do not predict such levels of variability [13]. Infinitesimal models of population genetics cannot explain the maintenance of high levels of phenotypic variation, in these models fluctuation selection predicts population crashes if variance is only explained by genotypes [20–22]. From an empirical point of view, estimates of heritability of functional traits and resulting expectations of trait shifts frequently mismatch with observed fluctuations in phenotypic traits of natural populations [13,23,24]. These challenges in explaining observed variability only by genotypes, environments and their interaction, lead us to the condition that non-genetic and non-environmental processes generate and contribute to the high levels of variation in phenotypes and life courses among individuals[11,12,14–17].

The fundamental question we address here concerns whether such non-genetic, non-environmental driven variation is truly neutral, selected for, or against. In the case of neutral variation, the follow up question would be, how is such neutral variation maintained [25]? Our approach here is not to decompose variance in genetic, environmental, phenotypic plastic (gene-by-environment), and neutral contributions to life course variability, as previously done for datasets that have the depth of information or by drawing on assumptions about partitioning [14,15,17]. Here, we aim at quantifying the selective forces on the processes that generate variation among life courses in relying on structured population theories [26]. We describe this approach in the following section starting with structured populations and associated life courses.

In any structured population, a life course of an individual can be described by a sequence of stages that ends with death [27]. These stage sequences, or life course trajectories, differ among individuals in length, i.e. age at death, and in the sequence and frequency of stages experienced. In many situations life of any individual starts in the same stage, the newborn stage. Thereafter life diversifies with increasing age, and the rate at which these sequences diversify with increasing length can be quantified by population entropy [4,26,28]. High entropy leads to highly diverse life courses in short times, and low entropy leads to few distinct life courses groups of individuals follow [28]. The life courses, i.e. the stage sequences, but also their diversification are determined by the stage transition rates[27]. In quantifying the effect of perturbations on each stage transition rate, i.e. each matrix element of the population matrix, we can evaluate the contributions of these perturbations to population entropy. In doing so, we quantify the contributions of each stage transition to the process of diversification of life courses[29]. Such estimation of the sensitivity of each transition rate to the population entropy does not reveal anything about fitness—λ, the rate at which a population grows [26,27].

Such a linkage to fitness can be acquired by the sensitivities of the transition rates to the population growth rate, λ. If one then correlates the sensitivity of the transition rate with respect to entropy and that to fitness, we acquire the link between life course diversification and selective forces (Fig. 1) [26]. To expand on this argument, if perturbations, i.e. sensitivities of a stage transition parameter with respect to entropy is correlated with the sensitivity of the same transition parameter with respect to the population growth rate (fitness λ), selection for diversification of life courses is favoured, whereas, if a negative correlation between these sensitivities occurs, selection against diversification is suggested, and if there is no correlation between the two sensitivities, the observed variability among life courses might be neutral. We motivate our interpretation with respect to selective forces on the idea that selection should act more strongly on stage-transitions that have higher sensitivities with respect to population growth, λ, and hence fitness [30]. To illustrate the concept, imagine a mutation that changes a vital rate (any fertility rate or stage transition rate), if this change in transition probabilities influences fitness, λ, more than changes in other vital rates, it should be under stronger selection than those vital rates that only have little influence on fitness.

**Fig. 1.**
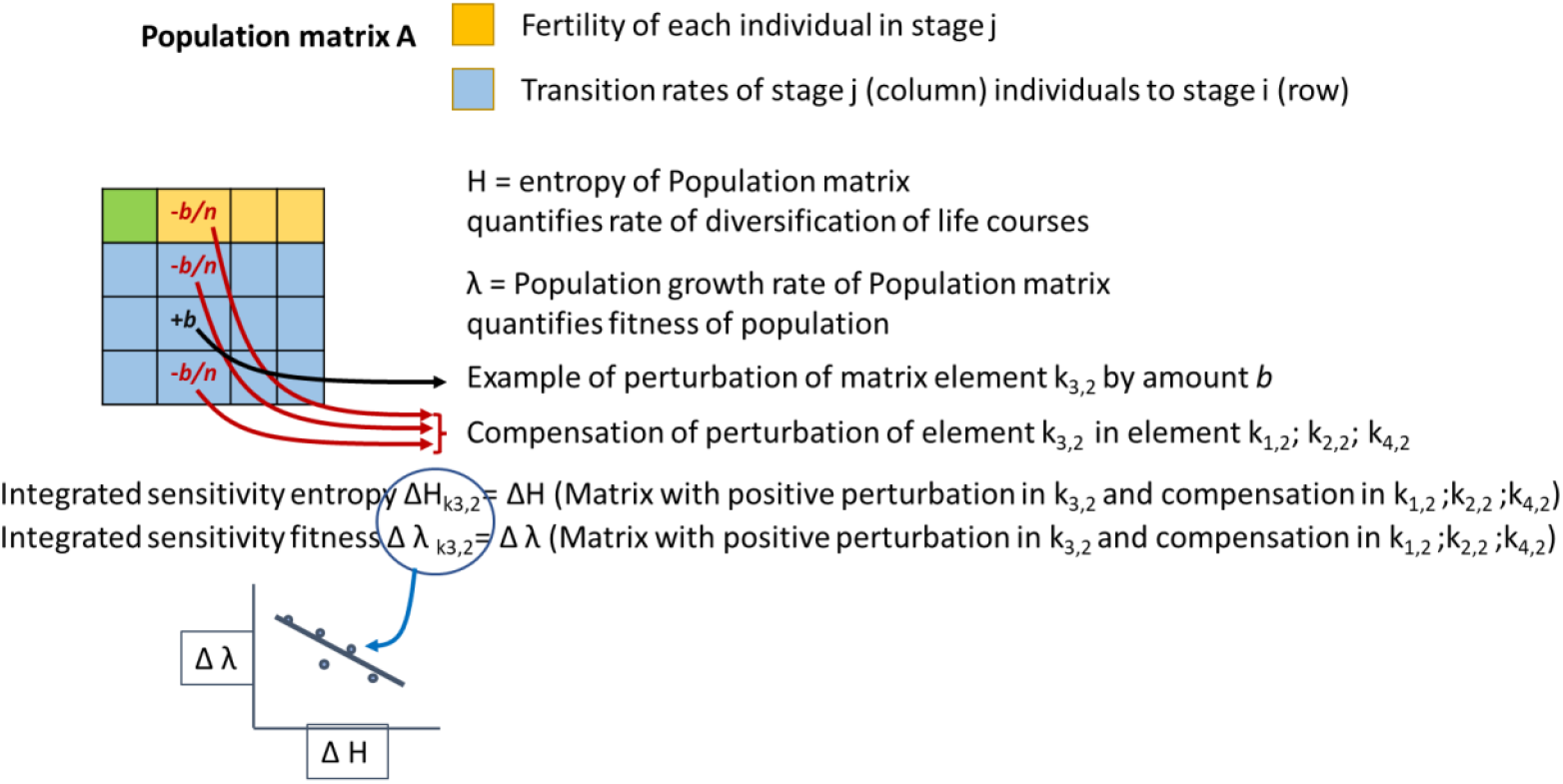
Sketch: for each population matrix, we estimated for each element (here exemplified by element k_3,2_) an integrated sensitivity with respect to entropy (ΔH_k3,2_) and with respect to fitness (Δ λ _k3,2_) by increasing (perturbing) element k_3,2_ by amount b and simultaneously reducing elements k_1,2_; k_2,2_; k_4,2_ by b/n with n=1-(Number of non-zero column elements). Such integrated sensitivities were then computed for each matrix element k_i,j_ and for both types of sensitivities. For each population matrix we fitted a linear model through data points based on these two types of sensitivities from each of the k_i,j_ elements. Each line in Fig. 2 corresponds to such a correlation model.

To evaluate how the diversity in life courses is selected upon—positively, negatively, or neutral—, we explore the correlation of the sensitivity with respect to entropy and the sensitivity with respect to population growth for a large variety of species and taxa for which population projection models have been collected within the COMADRE and COMPADRE data base [31]. We estimate for each transition rate of each population projection model the sensitivity with respect to entropy and population growth, then correlate these two sensitivities for each projection model, and compare these correlations across species, taxa, phyla, ontology, age, organism type and matrix dimension, for plants and animals. We find that both in plants and animals, substantial variation in the correlation between the two sensitivities among species exists, and we find a very weak or no overall correlation between sensitivities, suggesting close to neutral evolution of life course variability.

We also address a different question, whether populations or species with high rates of life course diversification, i.e. more diverse life courses, exhibit high fitness compared to those that are less diverse in their life courses. Such investigation might be understood in terms of adaptive niche differentiation or specialization[28]. Here, our findings suggest that matrixes with high rates of diversification (higher entropy) do not show increased or reduced fitness. Note, only a single entropy and a single population growth rate is calculated per matrix, while for each of the many non-zero matrix element sensitivities can be calculated. Overall populations that show greater diversity of life courses do not show increased or decreased population growth rates and selective forces seem not to increase or decrease life course variability.

## MATERIAL & METHODS

Out of the 3317 population matrixes in the COMADRE animal database and the 8708 matrixes in the COMPADRE plant database, we selected matrix models that were ergodic and irreducible (1350 and 5823 respectively). Of these, we selected only matrixes that had for each stage (each matrix column, Fig. 1) at least two non-zero elements; resulting in 37 matrixes on 11 animal species, and 2144 matrixes on 262 plant species. The extreme reduction in the animal matrix number reflects that many of these animal matrixes are sparse matrixes, for instance age-structured only (Leslie) population matrixes.

We limited the analysis to matrixes with at least two non-zero elements as to evaluate perturbations (sensitivities) that do not trade-off against survival, but against other stage transition or reproductive rates (Fig. 1). We call these sensitivities, that do not trade-off against survival, integrated sensitivities[26]. Each integrated sensitivity evaluates by how much a perturbation of amount *b*, in one focal matrix element *k*, influences population entropy, *H*, and population growth, λ, when simultaneously all other non-zero elements in the given stage (column) are reduced by *b/n*, with *n* equals the number of non-zero elements in a column minus the focal element. Note, integrated sensitivities can have negative and positive effects on entropy or λ.

Before we estimated the integrated sensitivities we transformed the absorbing population projection matrixes into Markov chains [32]. We then computed for each of the 41812 non-zero matrix elements their integrated sensitivities with respect to population entropy and population growth rate λ on the plant matrixes, and 602 non-zero elements of the animal matrixes. As the integrated sensitivities had very heavy tail distributions on both tails, we excluded extreme values that likely arose from biologically unrealistic matrix parameter entries or transition rates that were close to 1 or 0. We excluded extreme values of integrated sensitivities that exceeded three times the standard deviation for integrated sensitivities of entropy (13 animal matrix elements; 811 plant matrix elements) and values on integrated sensitivities of lambda that exceeded three times the standard deviation for the animal data (16), or 0.02 for the plant data (559), leaving 577 integrated sensitivities and 40640 integrated sensitivities respectively for the animal and plant data analysis (4 and 198 were outliers for both integrated sensitivities of respectively animal or plant data). Resulting distributions, after the outlier removal, remained heavy tailed.

For statistical testing, we fitted linear models (despite symmetrically long tails on both sides of the residual distribution) and used model comparison based on Akaike’s information criterion (AIC). We defined a difference in AIC>2 as substantial better support[33]. We evaluated the model fit and the assumptions using diagnostic plots.

For each matrix we also computed population matrix level entropy and population growth rate, λ, note there is one value of entropy and population growth for each matrix. We also correlated sensitivities with respect to entropy and those with respect to lambda for the 37 animal matrixes and 2144 plant matrixes against each other. Model comparisons were done using AIC’s[33].

## RESULTS

Across all animal species there is no evident correlation between the integrated sensitivities of entropy and those of lambda (Table 1: Model 1 [null model], vs Model2 [simple regression, slope -0.084], Fig. 2A), both models receive equal support. Hence, neither selection for nor against the diversity of life courses is observed in animals. For plants we find a weak positive correlation (Table 1: Model 1 vs Model2, Fig. 2B), though its effect size (slope 0.056) is small (compare with effect size of non-significant animal data). Hence, for plants selection tends to favour diverse life courses. This said, there is substantial variation in the correlation between the two integrated sensitivities among the species (Fig. 2; Table 1: Model 8 vs. Model 1,2,6,7), significant variation among matrixes (Fig. 2, Table 1: Model 4 vs Model 2, 3, 5), and significant variation among matrixes within species (Table 1: Model 4 vs. model 8). These findings suggest that selection differs among species, and matrixes within species, i.e. favouring diversity in life courses in some species while selecting against such diversity in others. Comparisons among matrixes for one specie show differences among populations, or the same population in different years. The results reveal that variation within species and among populations and years is also substantial. These patterns of substantial variance within and among species hold for animal and plant data.

**Table 1:** Model selection for animal and plant matrix data among competing models evaluating the correlation between integrated sensitivities with respect to entropy (response variable) and integrated sensitivities with respect to population growth lambda (explanatory variable) and various covariates.

|  |  | Animals |  |  | Plants |
| --- | --- | --- | --- | --- | --- |
| Model# | Parameters | DF | AIC | DF | AIC |
| 1 | Intercept only model | 2 | -7206.0 | 2 | -514160.2 |
| 2 | SensLambda | 3 | -7206.8 | 3 | -514477.0 |
| 3 | SensLambda+Matrix | 39 | -7169.5 | 2146 | -515011.4 |
| 4 | SensLambda*Matrix | 75 | **-7328.0** | **4289** | **-530900.5** |
| 5 | Matrix | 38 | -7166.0 | 2145 | -514832.1 |
| 6 | Species | 12 | -7203.3 | 263 | -517217.2 |
| 7 | SensLambda+Species | 13 | -7204.5 | 264 | -517418.6 |
| 8 | SensLambda*Species | 23 | **-7241.3** | 525 | -525791.6 |
| 9 | SensLambda*Species+Matrix | 49 | -7207.1 | **2407** | **-523644.7** |
| 10 | SensLambda*Age | 5 | -7205.2 | 5 | -514541.0 |
| 11 | SensLambda+Age | 4 | -7207.0 | 4 | -514484.8 |
| 12 | SensLambda*Matrix Dimension | 5 | -7204.4 | 5 | 514536.1 |
| 13 | SensLambda+ Matrix Dimension | 4 | -7206.3 | 4 | 514475.0 |
| 14 | SensLambda*Phylum | 9 | -7204.1 | 13 | -514793.7 |
| 15 | SensLambda+Phylum | 6 | -7201.6 | 8 | -514587.3 |
| 16 | SensLambda*Organism Type | 11 | -7202.1 | 21 | -515133.9 |
| 17 | SensLambda+ Organism Type | 7 | -7201.3 | 12 | -514491.7 |
| 18 | SensLambda*Ontogeny | 5 | -7202.8 | 5 | -514555.5 |
| 19 | SensLambda+ Ontogeny | 4 | -7204.8 | 4 | -514476.0 |

**Fig. 2:**
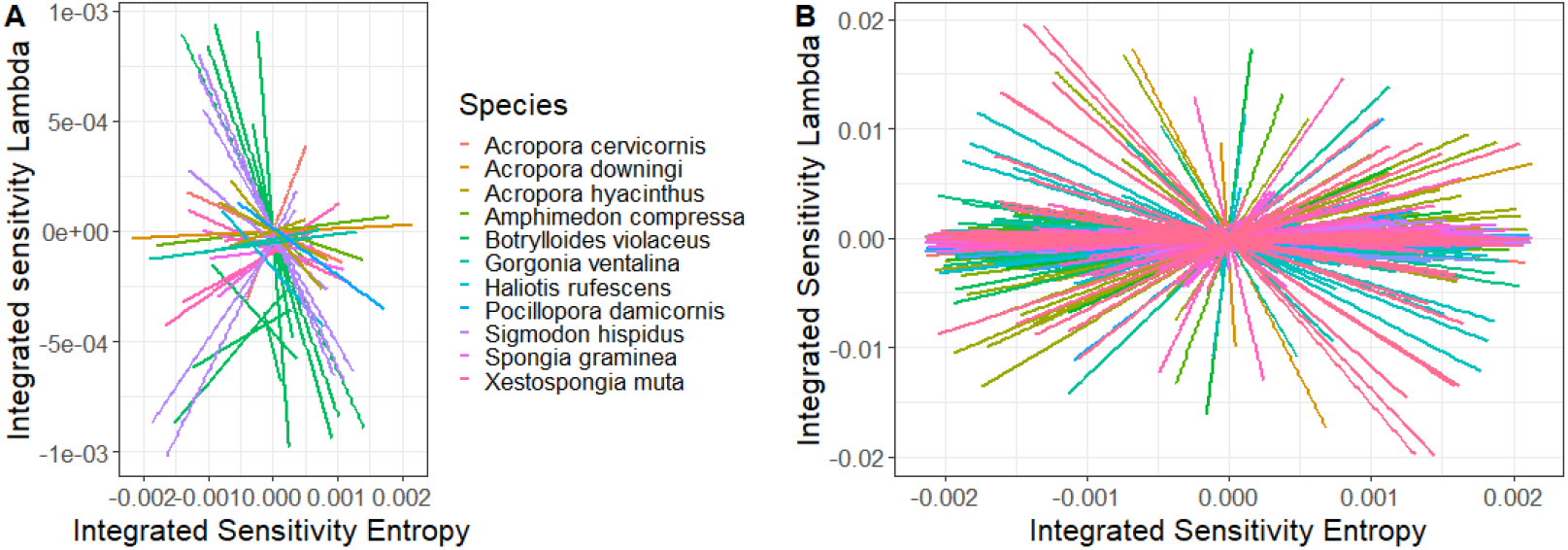
Correlating integrated sensitivities with respect to entropy and that with respect to lambda for animal populations (A) and plant populations (B). Each line fits the correlation for one population (one matrix model). Line colors reflect the different species as more than one matrix model can be fitted per species (e.g. different years, or populations). For the plant data (B) the number of species is too large to differentiate among the species. For better visibility CI (Confidence intervals) are not plotted.

Investigating other grouping variables in addition to the overall correlation among the two sensitivities, including age, matrix dimension, phylum, organism types, or ontogeny (Table 1, Fig. S1), shows that in animals these variables do not play an important role, while in plants they explain additional variability. Still, compared to the variance among species and within species, these grouping variables are of little importance. The number of stages per matrix (dimension) could potentially bias our findings. We found an interaction among matrixes of different dimensions and integrated sensitivity with respect to lambda for plant species, but there was no general trend with increasing matrix dimension, towards or against selection for variance in life courses, suggesting no systematic bias regarding matrix dimension (Fig. S1).

We further ask whether high or low diversity in life courses (population entropy) is associated with high or low fitness (population growth rates). Note here, we evaluate population entropy and lambda for the total population, i.e. one value for each matrix, not as above, a measure at the matrix element level (integrated sensitivities measures). Fig. 3 shows this relationship between entropy and lambda (see Table 2 for model comparison). We did not find any simple relationship between population entropy and fitness for animals (Table 2 Model 1 vs. 2, Fig. 3A), though for plants there was some tendency that matrixes with higher rates of diversification had lower fitness (Table 2 Model 1 vs. 2, Fig. 3B, slope - 0.42). One needs to caution these results as they are largely driven by extreme, biologically difficult to explain, high values of population growth rates (see also Fig. S2). Overall, significant variation exists between population entropy and population growth without clear correlation among the two variables. Matrix dimension explains some additional variance in the relationship between entropy and lambda, though species differences are much more important in explaining variance than matrix dimension.

**Fig. 3:**
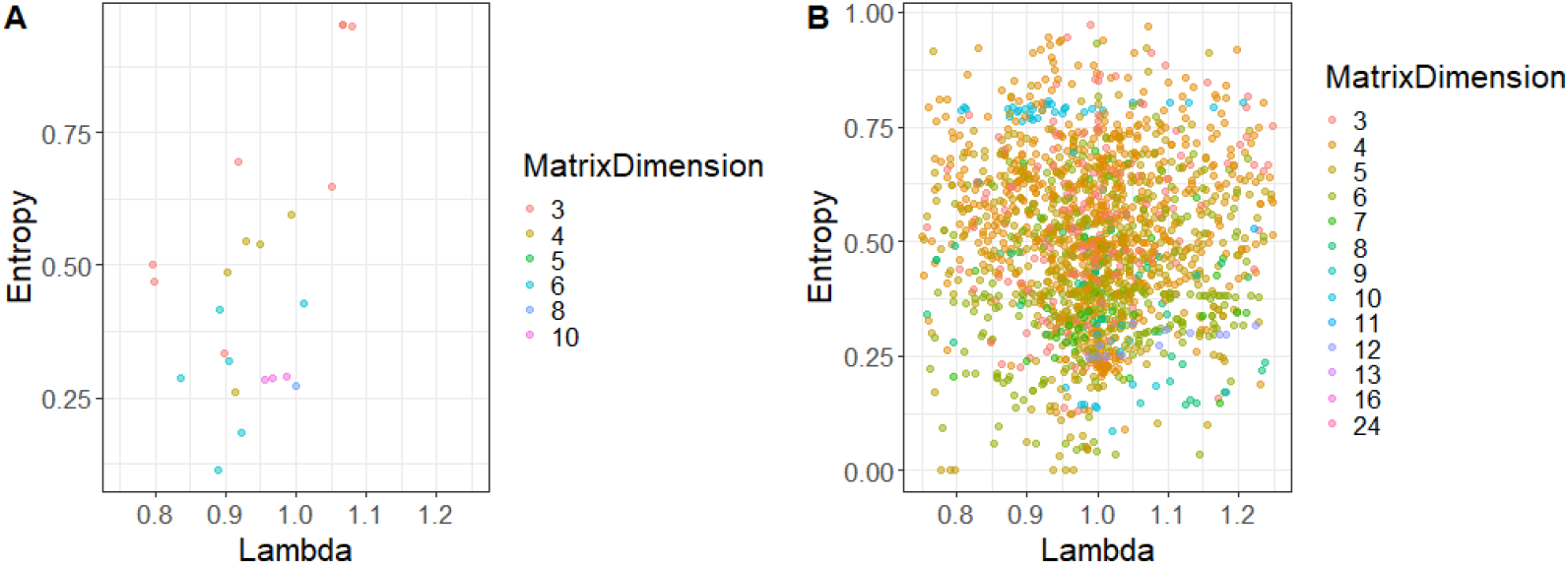
Relationship between population growth lambda (fitness) and population entropy, the rate of diversification, for animal (A) and plant (B) population models. Each data point represents one matrix model. Colors depict different dimensions of the matrix model. Populations that showed extremely low or high lambda are not plotted for better illustration. The full dataset, including the extreme values of lambda is plotted in Fig. S2 and the model selection of Table 2 is also based on the full data set.

**Table 2:** Model selection for animal and plant matrix data among competing models evaluating the correlation between population entropy (response variable) and population growth, lambda (explanatory variable), as well as matrix dimension and species comparison

|  |  |  | Animals |  |  |  | Plants |
| --- | --- | --- | --- | --- | --- | --- | --- |
| Model# | Parameters | Slope<br>SensLam. | DF | AIC | Slope<br>SensLam. | DF | AIC |
| 1 | Intercept only model |  | 2 | 50.19 |  | 2 | 2839.36 |
| 2 | SensEntr | 0.46 | 3 | 50.54 | -0.42 | 3 | 2779.30 |
| 3 | SensEntr+MatrixDim |  | 4 | 52.02 |  | 4 | 2780.43 |
| 4 | SensEntr*MatrixDim |  | 5 | **47.25** |  | **5** | **2754.89** |
| 5 | MatrixDim |  | 3 | 52.16 |  | 3 | 2835.28 |
| 6 | Species |  | 12 | 19.16 |  | 263 | 1625.00 |
| 7 | SensEntr+Species |  | 13 | **0.47** |  | 264 | 1624.83 |
| 8 | SensEntr*Species |  | 20 | 3.52 |  | **456** | **1583.70** |

## DISCUSSION

We show that across animal populations of various species no clear selective force acts towards or against increased or decreased diversification in life courses, whereas for plants an overall small positive slope indicates some level of selection favouring diversification in life courses. This latter finding in plants seem to contrast our finding that plant populations with high rates of life course diversification tend to have lower fitness than species or populations that show high rates of diversification. However, the two measures, the selective forces acting on the generating process of diversification in life courses, and the rate of diversification, reveal two different aspects. The integrated sensitivity analyses investigate selection on diversification processes within the population independent of the current rate of diversification the population exhibits [26], whereas population entropy quantifies this current rate of diversification [4]. The sensitivity analyses therefore focus on within population selective processes, whereas entropy and population growth are best used for among population comparison.

Our finding of substantial variation in selective forces on the generating processes, as well as substantial variation in the rate of diversification, might be of greater interest than the small positive selective trend favouring diversification for plant species. These substantial levels of variability might have three different biological origins or meanings: first, they might indicate substantial (developmental) noise that leads to the observed variability in life courses and selection for or against diversification in life courses[34], second, it might indicate fluctuating selection or high levels of phenotypic plasticity driven by variable environmental conditions[35,36], or third, it might indicate large numbers of distinct adaptive life courses that show similar fitness but might for instance fill different niches[28]. In quantitative genetic terms these options would relate to respectively, undetermined residual variation, gene by environmental variation, or additive variation.

If one assumes that noise explains the variability, it is suggested that selection might not act very strongly on this noise, as otherwise the variability should be selected against and variability should collapse[5–8]. Such neutral, or close to neutral, arguments have been used in the past to explain life course variability but are often met with scepticism[1]. Our results might indicate that selective forces on rates of diversification in life courses are not generally weak, but partly go in opposing directions, i.e. selecting for diversification in some populations or species and against in others, such conflicting findings are not uncommon in quantitative genetic studies[13,37,38].

If one assumes that fluctuating environments, or similar extrinsic variation, causes vital rates to differ among matrixes and leads to the highly diverse life courses[36,39], we might assume that a large fraction of variability would be explained by among matrix models *within* species, and less so *among* species. Model selection indicates that among species variation is substantially greater compared to variability among matrixes within species. Hence, variability among populations or time (years), or conditions (environments) within species contribute less to variability in life courses than variability among species. These arguments align with findings that phenotypic plasticity might not be in general adaptive [40]. The meta-analysis we did might not be ideal for such within species evaluation, as the average number of matrixes per species (3.4 for animals, and 8.2 for plants) are not very large, but our analysis still provides more general insights compared to studies focusing on single model species for which rich data exist[13].

If one assumes large number of adaptive life courses to drive life course diversity[28,41], we would be challenged to explain the strong selective patterns against diversification that is observed for some populations and species; though differences in population size might explain some of such variability. Under such assumption of distinct adaptive life courses, the optimal number of distinct life courses would need to differ substantially among species or populations. Also, from more detailed analyses of systems, certain life courses, or genotypes, that are commonly observed seem to have low fitness [13,14], suggesting that not all life course variability might be adaptive.

The three potential explanations that help to understand the selective forces on diversification of life courses are not mutually exclusive and we do not have means to quantify each contribution to the diversification using the meta-analysis data used in this study. More detailed studies that focus and explore selection on diversification could help to better understand the influence of these three factors[13]. Studies might include how genes (or gene knockouts) influence the rate of diversification, how experimental evolution studies in stochastic environments differing in amplitude and autocorrelation (noise color, wavelength) would lead to the evolution of different rates of diversification, or how “heritability” of distinct life course strategies potentially determine life course diversification under different environmental conditions. Quantitative genetics studies identified similar lack of understanding of maintenance and the evolution of variability[37,38], though with a focus on genetic explanations emphasizing mutation-selection balance being driven by few strongly deleterious mutations[38,42], or alternatively many polymorphic loci that maintain variability [37,43]. Such genetic variation interacts with neutral and non-genetically determined processes that influence evolutionary processes and the pace of evolution[1]. For that a purely quantitative genetic vision might be too short sighted. Generally, we believe only to be at the beginning to understand selection on processes that lead to the observed variability in life courses[13]. Increased interest in stochastic gene expression and its scaling and cascading effect across biological organization illustrate efforts towards such understanding[44,45].

---

**Fig. S1:**
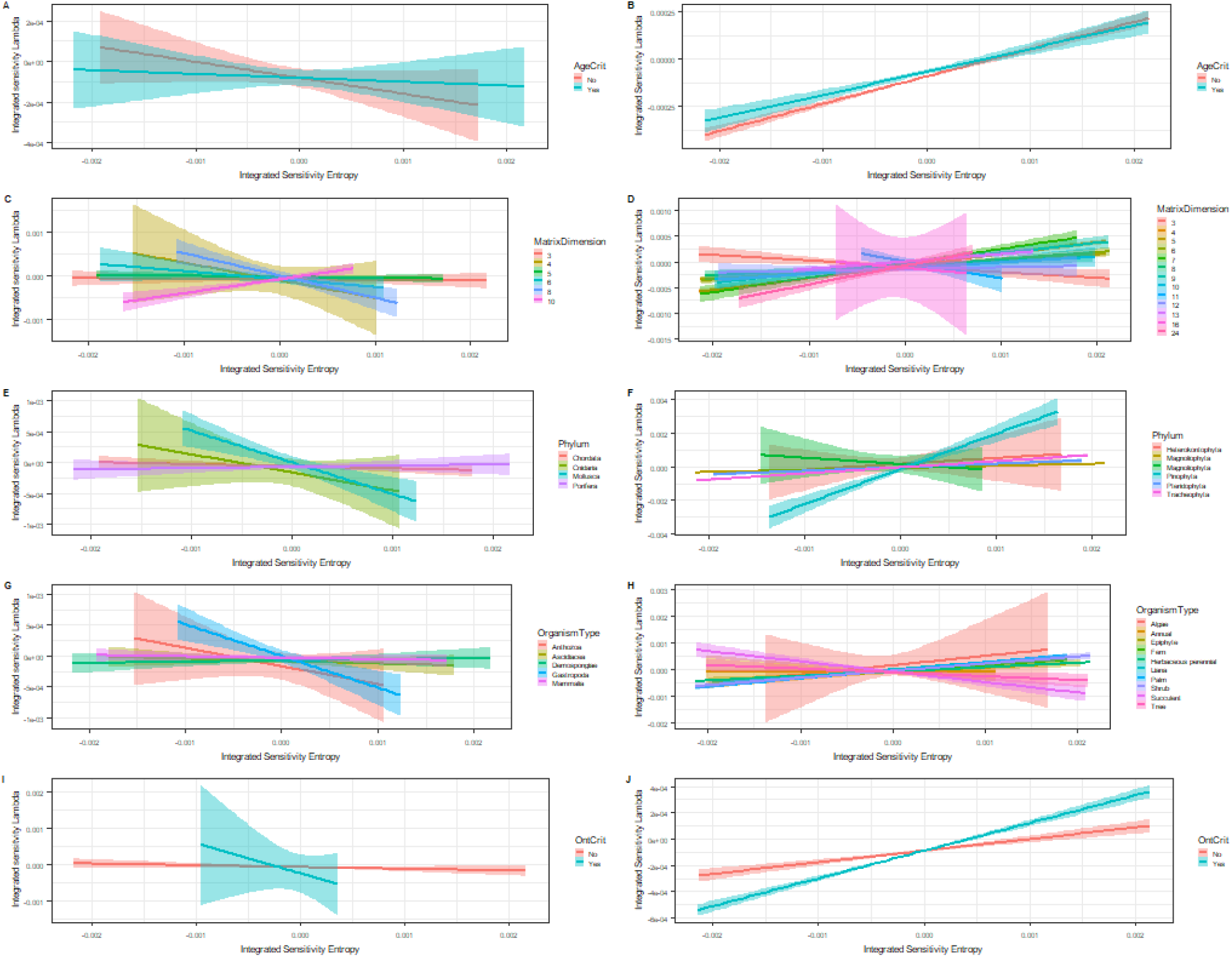
Correlation between integrated sensitivities with respect to entropy and integrated sensitivity with respect to lambda for animal populations (A, C, E, G,I) and plant populations (B,D,F,H,J). Grouping variables include whether the matrixes are based on age or not (A&B), their dimension of the matrix [number of stages] (C & D), their Phylum (E &F), their Organism type (G & H), or whether they are based on ontogeny or not (I & J). Each line fits the correlation for one of the grouping variables with the 95% CI. Note, we plotted for better visualization the matrix dimension (C &D), one regression for each dimension, though for the statistical models (Table 1) matrix dimension was used as a continuous variable.

**Fig. S2:**
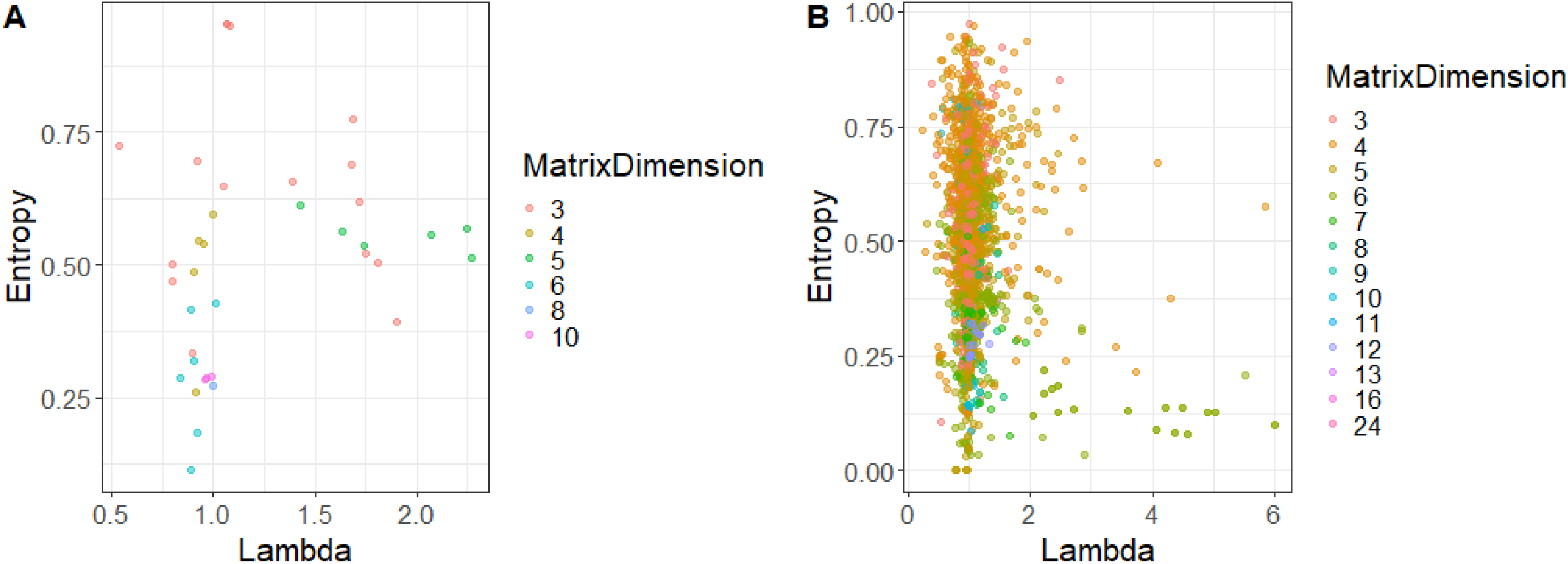
Relationship between population growth lambda (fitness) and population entropy, the rate of diversification, for animal (A) and plant (B) population models. Each data point represents one matrix model. Colors depict different dimensions of the matrix model. Here the full dataset, including the extreme values of lambda is plotted.

